# An epidemiological description of rabies outbreaks in Zambia from 2013 to 2023

**DOI:** 10.1101/2024.07.16.603659

**Authors:** Humphrey Banda, Amos Hamukale, Chitwambi Makungu, Masuzyo Ngoma, Ricky Chazya, Liywalii Mataa, Chilufya Mulenga, James Blazer Banda, Walter Muleya, Cephas Sialubanje, Dabwitso Banda, Nyambe Sinyange, Wezi Kachinda

## Abstract

**Background:** In Zambia, a dog bite is the most common rabies exposure. There is limited information on rabies outbreaks in jackals in Zambia. We investigated all rabies cases in Zambia and narrowed down on jackal bites in Mungwi district to determine a temporal trend and buffer zones for rabies.

**Methods:** The risk of exposure included all humans and animals bitten by jackals, data were collected by interviewing jackal bite victims and collecting brain specimen samples from animals. Risk of spread was determined by estimating the distance jackals strayed into human population and the Rabies Vaccination Coverage (RVC) in the district. The RVC was determined by dividing vaccinated dogs by dog population. The incidence rate (IR) and a Sen’s slope was used to determine a trend of RVC and the rabies cases in Zambia. QGIS was used to produce a heatmap and rabies risk zones. Direct Fluorescent Antibody technique was used to test for rabies.

**Results:** Zambia had recorded a total of 224 rabies cases with a mean (SD) of 22.4 (16) rabies cases and a positivity rate of 65.4% (25.7) per year from 2013 to 2022. Lusaka and Copperbelt provinces recorded the highest rabies cases. The rabies incidence rate was 2.3 rabies cases per 10,000 dogs in Zambia. The cases had reduced from 2013 to 2021 with a Sen’s slope of -0.74 (p = 0.02). At least 227 animal bites were recorded (two jackals and 225 dog bites) in Mungwi District. 94% (213) of the victims needed Post Exposure Prophylaxis. The victim’s median age was 20 (Interquartile range=12-38) years. The accumulative RVC from 2018-2022 (611/7777), was 0.11 (95% CI: −0.4-0.7, p=0.46) annual slope increase with 1.60% mean (Standard deviation = 0.34). One rabies heatmap and three risk-level zones (low, medium and high) were produced, and two jackals and one goat specimens were positive.

**Conclusions:** A risk-based surveillance and enhanced vaccination of dogs in high-risk areas through a one health approach is critical for rabies control in Zambia.

## Background

Rabies is a public health concern and a viral zoonotic disease (1). It is an RNA lyssavirus of the Rhabdoviridae family (2) and can be transmitted between all warm-blooded species including man. Studies have shown that several domestic and wild animals such as dogs, cats, livestock animals, wolves, foxes, jackals, and bats are susceptible to rabies virus and can transmit the disease to humans via bites or scratches (2). Human rabies is mostly due to dog-transmitted rabies virus through animal bites (3).

Rabies disease is endemic on all continents except Antarctica (4) and it causes approximately 59,000 human deaths every year, this translates to one person dying every 10 minutes (5). This disease is most prevalent in low and middle-income countries with poor or lack of management and control strategies (6).

Rabies occurrence in Zambia has been experienced sporadically in various parts of the country (7,8). The country reported 1935 rabies cases of which 1004 were confirmed (8), and 463 rabies cases from 2004 – 2014 (7). These studies reported that Lusaka province had more confirmed cases among all provinces (7,8). In Zambia, rabies infects a wide range of species such as dogs, cats, cattle, goats, sheep, and humans; also wild animals such as jackals, foxes, monkeys, and baboons (7,8).

The World Health Organization (WHO) recommends mass dog vaccination campaigns of 70% vaccine coverage and strict dog population control via restricted breeding, restricted movements, and stray dog cropping (9). Rabies post-exposure prophylaxis (PEP) prevents rabies in humans exposed to the rabies virus (10,11). Thus, PEP is the cornerstone for rabies prevention in humans, and it is against this background that it is recommended for all persons that have been or are suspected to have been exposed to the rabies virus. Despite rabies being a preventable disease with available effective vaccines, most countries are still experiencing outbreaks for quite a long time due to low vaccine coverage and irresponsible dog ownership (9). The extent of the outbreak and risk exposure to rabies in Mungwi District has not been established and there is limited information on the epidemiological trends of animal attack cases. We investigated to determine the proportion of animal bites, dog vaccinations and possible rabies exposure in the communities of Mungwi District (January-2018 to February-2023). The investigation provided recommendations on rabies prevention and control strategies.

## Methodology

### Study design

This was a retrospective and cross-sectional study design, analyzing primary and secondary data collected from the outbreak investigation and the National Livestock Epidemiological Information Centre (NALEIC) under the Department of Veterinary Services (DVS) and Central Veterinary Research Institute (CVRI) in Zambia.

### Study setting

The outbreak investigation included secondary data of all rabies cases found in the NALEIC and CVRI databases from January 2013 to December 2022. In addition, all dog rabies vaccinations and animal attack cases reported in Mungwi District in Zambia from January 2018 to December 2022. These cases were reported from veterinary camps. A veterinary camp is a subdivision of the district, which is manned by a Veterinary Assistant, who in turn reports to the District Veterinary Officer (DVO). The cases came from all the 10 provinces of Zambia (Figure 1).

**Figure 1.**
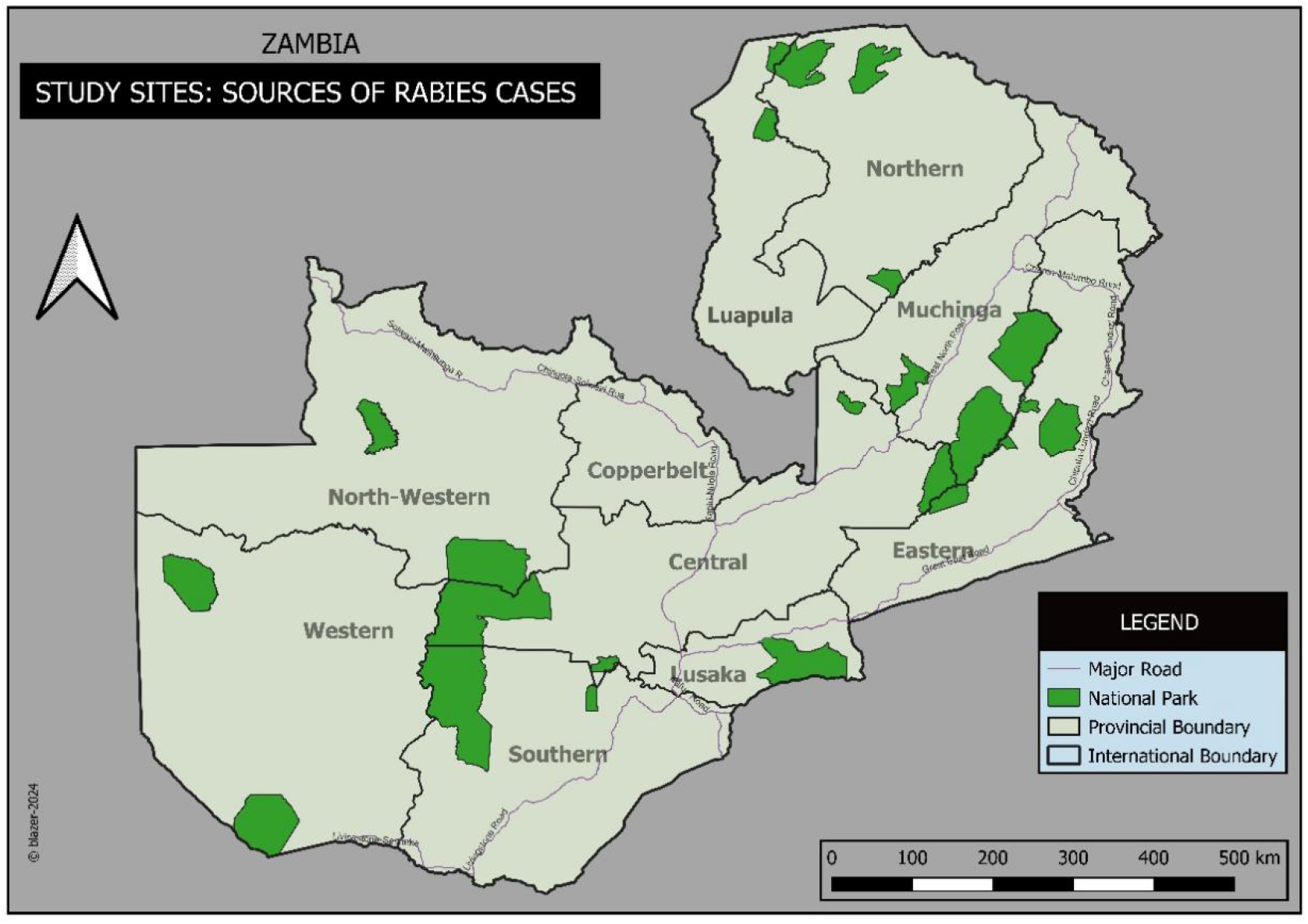
Study sites: Sources of rabies cases in the Provinces of Zambia from 2013 to 2022

### Sampling strategy

We collected all rabies vaccination and animal bites records at NALEIC and Mungwi DVO’s office (2018 - 2022), and all cases of rabies in animals in Zambia (2013 to 2022) from NALEIC and CVRI. Additionally, participants in the study included jackal bite victims and all dog bite cases reported within the affected areas in Mungwi District.

### Case definition

#### Rabies infection

##### Suspected case

A person or animal presenting with an acute neurological syndrome (encephalitis) dominated by forms of hyperactivity (furious rabies) or paralytic syndromes (dumb rabies) progressing towards coma and death, usually by respiratory failure, within 7-10 days after the first symptom and lives within the affected area of Mungwi District.

**A probable case** is a suspected case with a likely exposure to a confirmed rabid animal.

**A confirmed case** is a person or any animal with laboratory evidence of rabies infection by detection of either one of the following:

a. rabies virus nucleic acid by RT-PCR on saliva, skin biopsy or cerebrospinal fluid (CSF)
b. Anti-rabies antibodies in CSF (ante-mortem).
c. Rabies virus antigen in brain tissue by fluorescent antibody testing (FAT) or rabies virus nucleic acid in skin biopsy.

#### Animal bites and rabies exposures

According to the World Health Organization, the following are categories are defined as cases that require Post Exposure Prophylaxis (PEP):

**Category I**: Touching or feeding animals, animal licks on intact skin (no exposure).

**Category II**: Nibbling of uncovered skin, minor scratches or abrasions without bleeding (exposure).

**Category III**: Single or multiple transdermal bites or scratches, contamination of mucous membrane or broken skin with saliva from animal licks, exposures due to direct contact with bats (severe exposure).

#### Data collection

All rabies cases from 2013 to 2022 were extracted from NALEIC and CVRI databases using a data collection tool in excel. We interviewed jackal bite victims and key informants using an interview guide of open-ended questions focusing on rabies risk exposure and occupational hazards. The rabies vaccination and animal bites records were collected from the district monthly NALEIC reports. The coordinates for the jackal habitat and bite incidences were collected using a GARMIN eTrex® 10 GPS gadget. The dog populations were collected from the 2017/2018 Livestock and Aquaculture Census and the 2022 Livestock Survey report.

### Specimen collection

The specimens were collected by the response team at the provincial veterinary office for Northern Province.

### Data analysis

We calculated the proportions of demographic, risk exposure, and animal rabies vaccination data. Rabies Vaccination Coverage (RVC) was calculated as a proportion of dogs vaccinated in the District from January 2017 to December 2022, over the estimated dog population in the district. A rabies temporal trend from 2013 to 2022 in Zambia, was analyzed using a Sen’s slope test in r statistical software. The coordinates were used to generate a map showing the buffers zones for rabies exposure using Quantum Geographic Information System (QGIS). The incidence rate was calculated using the following formula:

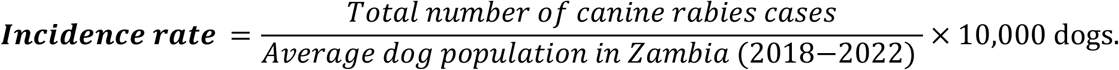

## Results

### Rabies temporal trend in Zambia

Zambia had recorded a total of 224 rabies cases with a mean (SD) of 22.4 (16) rabies cases and a positivity rate of 65.4% (25.7) per year from 2013 to 2022. Lusaka and Copperbelt provinces recorded 17% each and were the highest proportion of rabies cases recorded among provinces of Zambia. The rabies incidence rate was 2.3 rabies cases per 10,000 dogs in Zambia. Over 78% (176/224) of affected animal types were dogs and followed by cattle at only 8% (18/224). The cases had reduced from 2013 to 2021 with a Sen’s slope of -0.74 (p = 0.02) and then a sudden peak in 2022 suggesting a decreasing trend in reported rabies cases and the positivity rate in Zambia (Figure.2).

**Figure 2.**
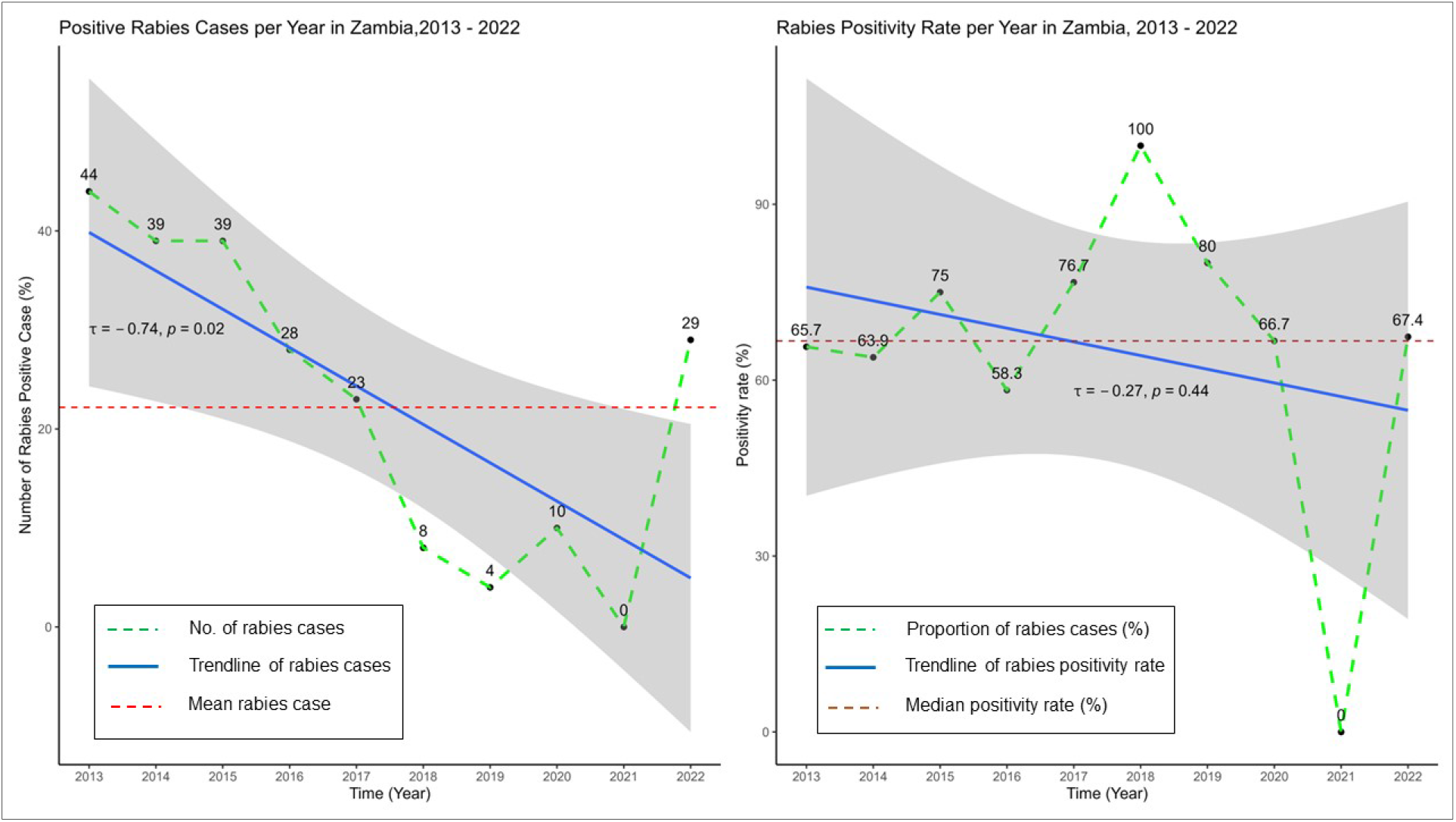
A temporal trend of rabies cases and the positivity rate in Zambia, 2013 – 2022

The rabies heatmap shows that Lusaka, Copperbelt and Eastern provinces are the most affected. Medium affected provinces are Central, Luapula and the northern part of Muchinga with the bordering districts of Northern Province. Also, some parts of Chirundu, Chikankata, Mazabuka, Monze and Siavonga districts of Southern Province. This also highlights that the fact that rabies is endemic in Zambia all provinces have at least reported a rabies case (Figure 3).

**Figure 3.**
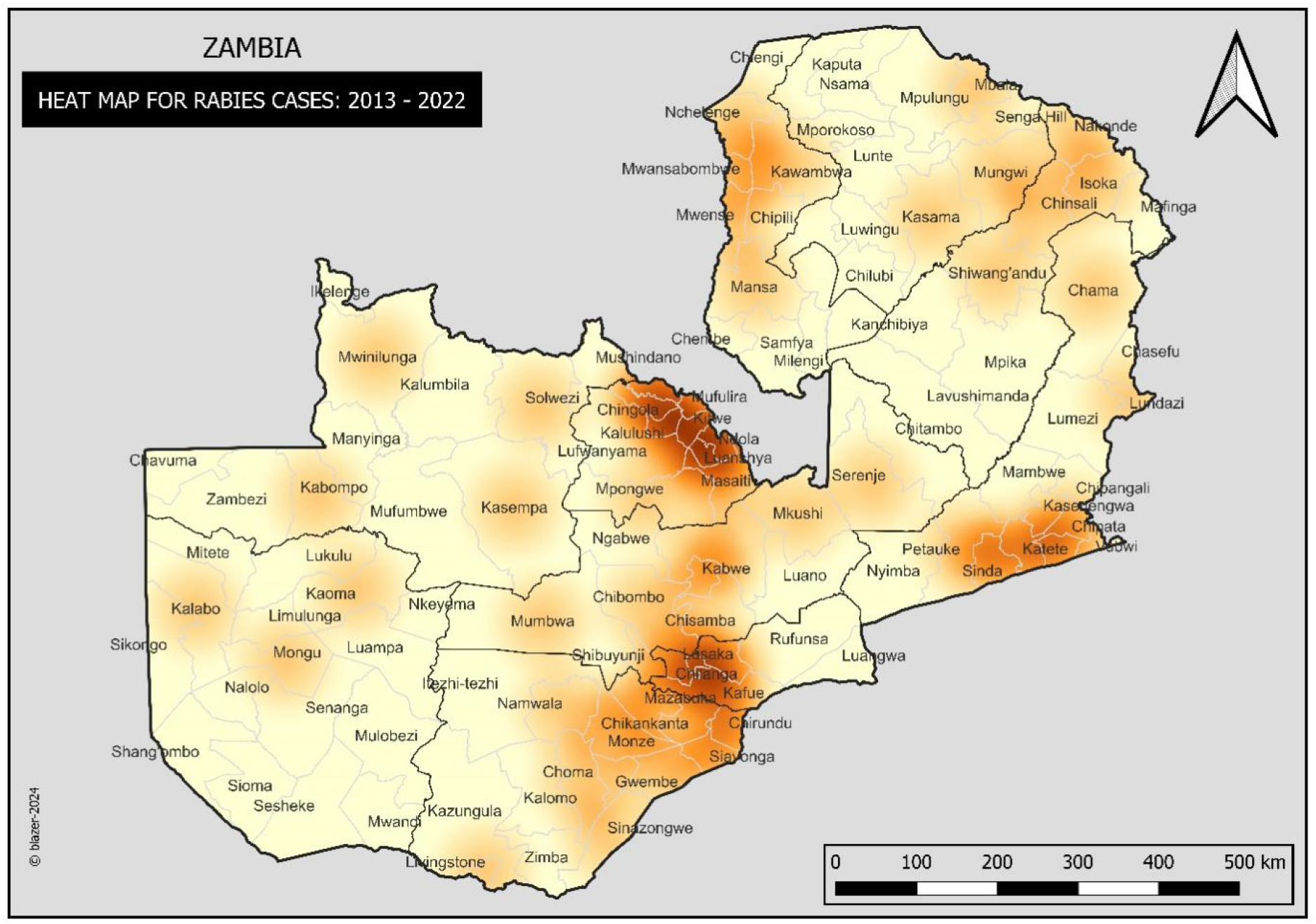
Rabies heatmap in Zambia from 2013 to 2022

**Figure 4.**
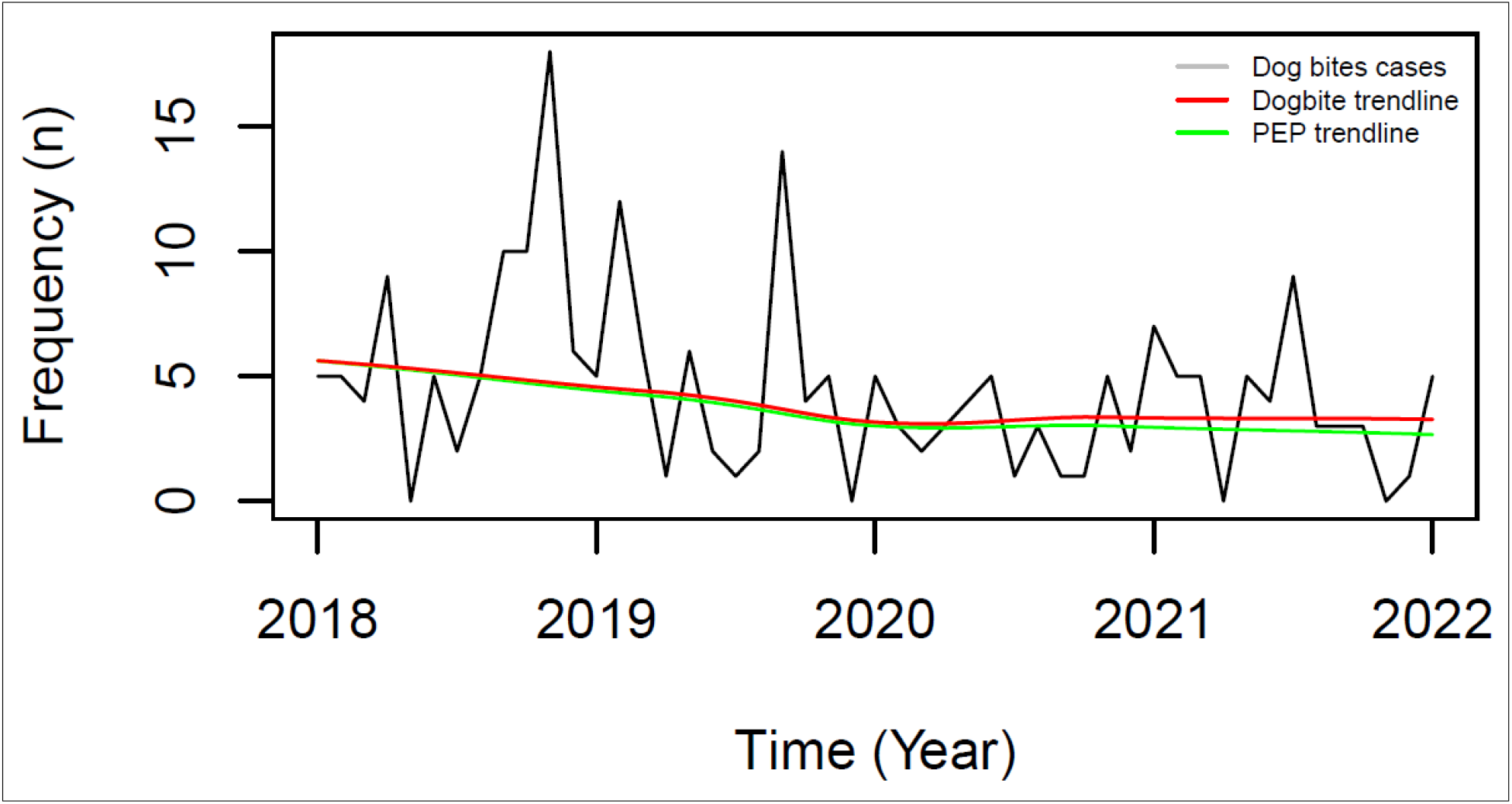
Animal bites cases by month and year in Mungwi District, 2018 – 2022

### Rabies temporal trend in Mungwi District

In the period between January 2022 to February 2023, Mungwi district recorded 48 (46 dog bites and 2 jackal bites) animal bites cases. The median age of the victims was 20 years with an IQR of 12-38 years. Out of the 48 animal bite cases, 81% (39) were recommended for PEP (Table 1).

**Table 1.** Animal bite cases in Mungwi District, Jan 2022 – Feb 2023

| Characteristic | Non-PEP | PEP needed | Totals |
| --- | --- | --- | --- |
| Animal bites | N = 9 (19%) | N = 39 (81%) | N = 48 (100%) |
| Age (years) | 16 (12, 36) | 21 (12, 37) | 20 (12, 38) |
| Unknown (age) | 2 | 0 | 2 |

Period
| Characteristic | Non-PEP | PEP needed | Totals |
| --- | --- | --- | --- |
| Jan – Dec 2022 | 8 (21%) | 30 (79%) | 38 (100%) |
| Jan – Feb 2023 | 1 (10%) | 9 (90%) | 10 (100%) |

A decreasing trend was noted in animal bite cases in Mungwi from 2018 to 2022. The cases had reduced from 2018 to 2020 with a Sen’s slope of -0.05 (95% CI: −0.13 to 0 and p = 0.03) and then steadily stabilized up to 2022 (Figure.1). Equally a decreasing trend of animal victims who were recommended for PEP was reducing with a slope of -0.07 (95% CI: −0.13 to 0 and p ≤ 0.009).

### Rabies vaccination coverage in Mungwi District

The study revealed a critically low rabies vaccination coverage (RVC) in Mungwi district, with an accumulative coverage of only 0.11 (95% CI: −0.4-0.7, p=0.46) from 2018-2022 (Figure 5), and a mean annual increase of 1.60%. This inadequate vaccination coverage correlates with the high demand for post-exposure prophylaxis (PEP), as 81% of bite victims required PEP.

**Figure 5.**
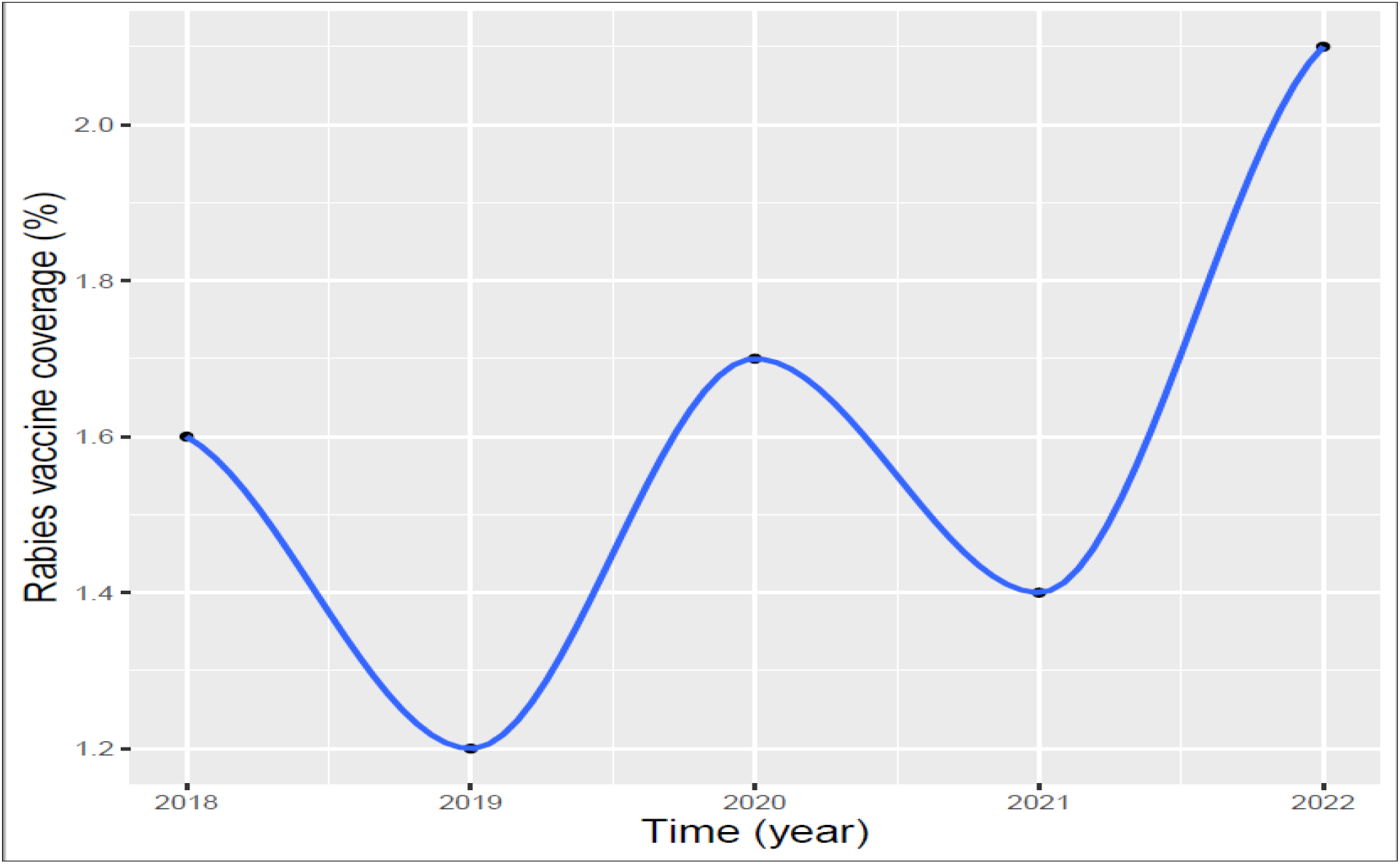
Rabies vaccine coverage in Mungwi District, 2018 - 2022

### Rabies outbreak in Mungwi District

There were two women aged 22 and 42 years respectively, two dogs and three goats that were bitten by two different side-strapped jackals in Mungwi District of Northern Province in February 2023 (Figure 6). The first woman was eight months pregnant and bitten on the umbilicus region, while the other was bitten on the left foot and right index finger. Both victims were given antrabies therapy (Post Exposure Prophylaxis). After 21 days of rabies exposure, one of the goats showed rabies clinical signs before it were killed. Out of the four samples submitted to the Central Veterinary and Research Institute (CVRI) laboratory in Lusaka, three (Two jackals and one goat) tested positive for rabies.

**Figure 6.**
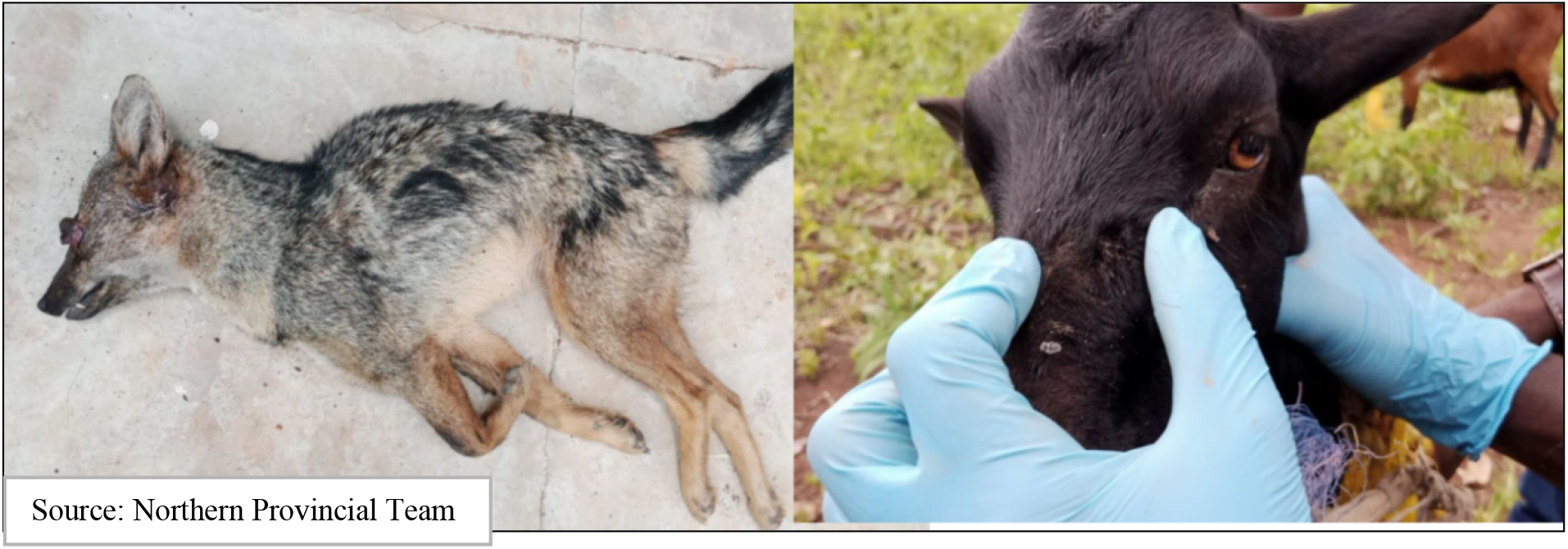
The side-stripped jackal and the goat in Mungwi District, February 2023

### Rabies transmission buffer zones in Mungwi District investigation

The study identified specific instances of jackal bites, which included two women, two dogs, and three goats. The distances between the jackal habitat and the bite incidents were 19 km and 7 km respectively. Three (Low, medium, and high risk) rabies possible risk transmission buffer zones were identified in the outbreak area in Mulambe and Mabula villages of Mungwi District (Figure 7).

**Figure 7.**
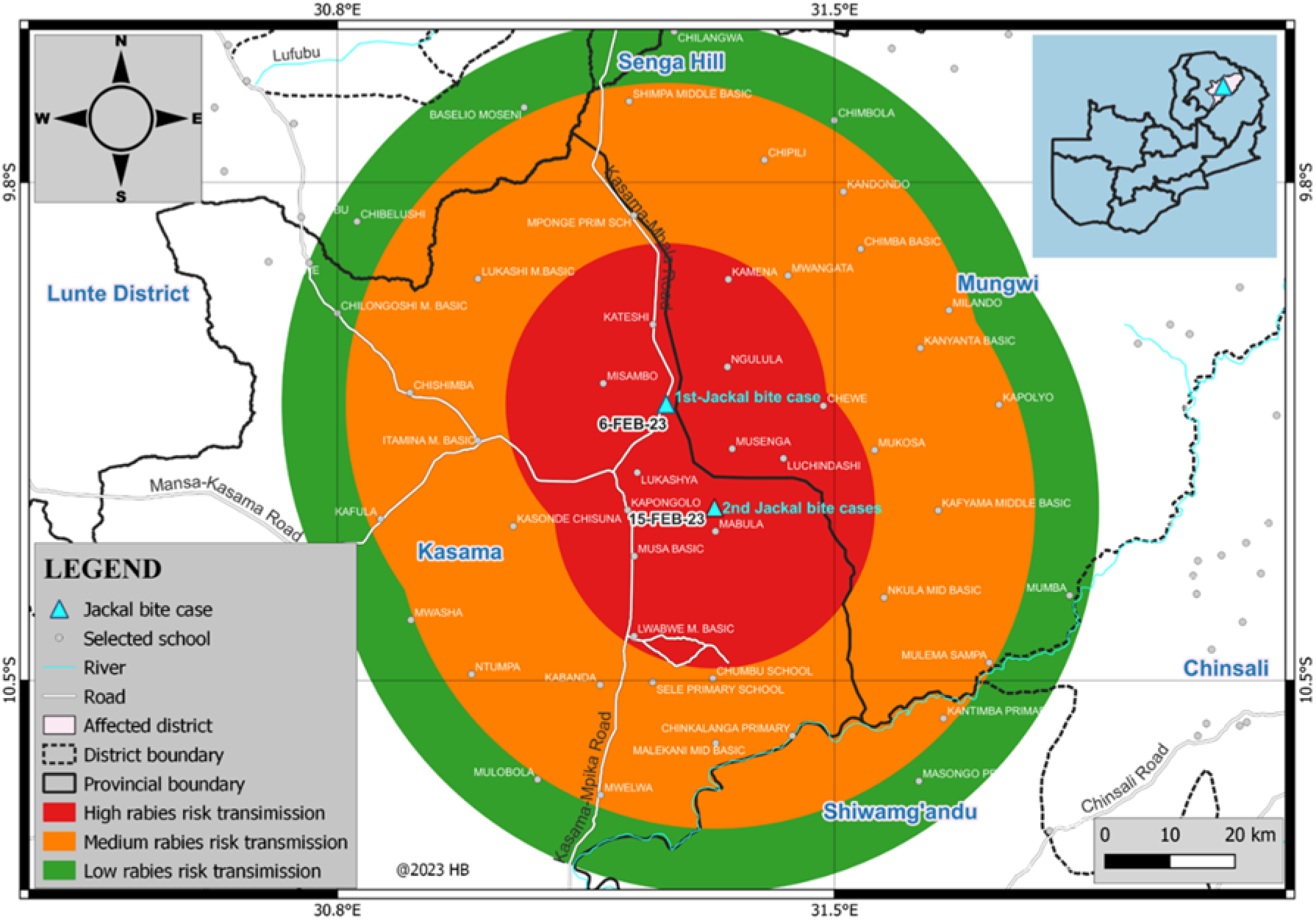
Rabies buffer zones in Mulambe and Mabula villages of Mungwi District, 2023.

## Discussion

### Rabies in Zambia

The study results confirm that rabies remains a public health concern in Zambia, despite it been rare, rabies is endemic in all provinces. This finding coincides with earlier studies conducted in Zambia (7,8). Compared to the period 1984–2004 and 2004 – 2014 (7,8), this study indicates a decreasing reported animal rabies cases in Zambia. This can be attributed to the realization of the public health impact of the disease by the authorities and the implementation of various interventions such as rabies vaccination and education programs in Lusaka Province. The other contributing factors could be probably under reporting which is a problem in most low income countries like Zambia (12). In most rural and remote areas, communities would kill and suspected rabid dogs without reporting to the veterinary offices and in some cases sample courier system does not exist and places far from Lusaka would only submit rabies suspected samples only if they have opportunities of a vehicle destined for Lusaka. At times, the DVOs do not even have postmortem kits which would force them to submit the whole carcass or the head of the suspected animal.

### Animal bites and Post Exposure Prophylaxis

Rabies is primarily transmitted to humans through the bite of an infected animal, with dogs being the main driver of rabies. In Zambia, dog bites are the most common route of rabies transmission. There is a notable discrepancy between the number of dog bite cases recorded by hospitals or clinics and the Veterinary Office. This discrepancy suggests that many dog bite cases may not be reported to the Veterinary Office, which could lead to an underestimation of the true situation in the case of Mungwi District. Despite the discrepancy, a reduction in dog bite cases over time was observed. However, a significant percentage of dogs involved in these cases remain unvaccinated, indicating a high demand for PEP (13).This is consistent with other studies in Zambia which found that the distribution of unvaccinated dogs in rural areas than urban or peri-urban areas (14). PEP is a treatment that can prevent rabies if administered promptly after a bite from an infected animal or in most cases unvaccinated dog. However, PEP is not always available in the district and when available it is only stocked in the district hospital which is in the urban area. The victims have the added costs of transportation which significantly may reduce adherence to PEP treatment (14). These additional costs can be reduced by vaccinating dogs against rabies. Besides, PEP has been proven to be more expensive than dog vaccination in terms of rabies control and may not be cost effective in the long run (13).

The other contributing factor to the low rabies dog vaccination coverage is that despite Zambia having an official control strategy on rabies and vaccination been mandatory, this disease is classified as a management disease which by definition entails that the pet owners bears all the cost of vaccination (15–17). The government we will only come in with free vaccinations during an outbreak. The high cost of rabies vaccination is another barrier which was identified in a study in Nyimba District of Zambia (14)

### Rabies in Wild Animals

Despite the limited understanding about rabies in wild animals in Zambia, occasional outbreaks of rabies have been reported in national parks and interface areas, suggesting that the virus may be present in wild animal populations. Domestic dogs are believed to be the primary transmitters of the rabies virus to humans and wild animals in Zambia, highlighting the importance of vaccinating dogs to prevent the spread of the disease. Nevertheless, other studies have found the clustering of rabies virus strains from jackals with those from domestic animals provides evidence of similar virus strains circulating in both wildlife and domestic animals, and that the jackals might be potential reservoirs of rabies virus in Zambia (18).

### Use of QGIS Mapping in Rabies Outbreak Response

John Snow famously demonstrated the power of plotting the spatial locations of individuals affected in an outbreak (19). His spatial analysis of cholera cases in London in 1854 showed a clear pattern that implicated a water pump as the likely source of infection. Nevertheless, epidemiological investigations of outbreaks have tended to focus more on analysis of person or animal and time than of place (20). Geographic information systems (GIS) have increased the availability and range of tools that can be used to analyze outbreaks (21). A GIS is a database designed to handle geographically referenced information complemented by software tools for the input, management, analysis and display of data. GIS are used widely in epidemiology and the simplest application in an outbreak investigation is to create maps displaying the relative locations of cases, potential sources and/or risk factors (20–22). Maps can help identify areas with high population density of unvaccinated dogs, allowing for targeted vaccination campaigns. Despite the potential benefits, spatial methods are underutilized and used in only 0.4% of all published outbreak investigations (21). mapping has not been extensively utilized in Zambia during rabies outbreak responses. Our study demonstrated the importance of mapping the outbreak areas and the creation of buffer zones for estimating the extent of the problem and conducting a targeted vaccinations and surveillance. This requires strengthening the capacities of the Department of Veterinary Services at district in cartography and outbreak investigation for effective response and better control of rabies outbreaks.

## Conclusions

A risk-based surveillance and enhanced vaccination of dogs in high-risk areas is critical for rabies control in Zambia. This requires strengthening the capacities of Department of Veterinary Services at district level in diseases cartograph, prevention and control through enhanced one health collaborative approach. A multidisciplinary approach in collaboration with all the line Ministries and public health institutes working together with local communities on rabies campaigns and strengthening of public awareness and education such as responsible pet ownership, dog bite prevention and rabies vaccinations will play a pivotal role in rabies control and elimination. There is need for extensive research on rabies in wildlife and the development of rabies risk maps for Zambia.

## Acknowledgements

We would like to acknowledge the support from Zambia National Public Health (ZNPHI), Zambia Field Epidemiology Training Program and Ministry of Fisheries and Livestock - National Livestock Epidemiology Information Centre (NALEIC), the Provincial Veterinary Office – Kasama, and the District Veterinary Office – Mungwi.

## Notes

### Competing Interest Statement

The authors have declared no competing interest.

